# No preference for performance: Host plant preference, offspring performance and host plant distribution in the butterfly *Aricia artaxerxes*

**DOI:** 10.64898/2026.03.02.708994

**Authors:** Vanda Larsson Åberg, Jesper Boman, Niclas Backström, Martin I. Lind

## Abstract

The connection between female host plant preference and offspring performance is important for understanding how relationships between plants and phytophagous insects have evolved. According to the preference-performance hypothesis, female insects should evolve to oviposit on host plants on which offspring performance is the highest. Here, we examined the preference-performance hypothesis in the northern brown argus (*Aricia artaxerxes*) butterfly in the province of Uppland, Sweden, by comparing female host plant preference and larval growth between the host plant species wood cranesbill (*Geranium sylvaticum*) and bloody cranesbill (*G. sanguineum*). We also investigated if host plant preference in *A. artaxerxes* was related to the geographic distribution of *A. artaxerxes* and its host plants in the province Uppland. We found that the *A. artaxerxes* females, contrary to the preference-performance hypothesis, preferred ovipositing on *G. sylvaticum*, even though larvae feeding on *G. sylvaticum* were slightly smaller than those feeding on *G. sanguineum*. Since *G. sylvaticum* is more abundant and probably more utilized than *G. sanguineum* in Uppland, an explanation for this negative preference-performance connection may be that there are advantages associated with utilizing a more common host plant species, even though larvae feeding on this plant show reduced growth rates. Overall, the results show that factors other than offspring performance, such as geographic distribution, may influence female host plant preference in *A. artaxerxes*.

## Introduction

Coevolution is the reciprocal evolutionary change occurring between interacting species through natural selection (Thompson 1989). These interacting species can be mutualists, such as flowering plants and pollinators (Bronstein *et al*. 2006), or antagonists, such as competitors (Connell 1980), predators and prey (Abrams 2000) and parasites and hosts (Anderson & May 1982). One example of antagonistic coevolution is that of plants and phytophagous (plant-eating) insects (Ehrlich & Raven 1964). Together, plants and insects compose more than half of the world’s described species (Futuyma & Agrawal 2009, IUCN 2025). However, phytophagous insect species, generally only utilize a restricted number of host plant species (Wiklund 1975, Bernays & Graham 1988, Thompson 1993). These host plant associations are shaped by plant secondary metabolites involved in the defense against herbivores (Ehrlich & Raven 1964, Bennett & Wallsgrove 1994). According to Ehrlich and Raven’s (1964) *escape-and-radiate coevolution* hypothesis, plants evolve new secondary metabolites that enable them to escape from herbivores and undergo radiation. When phytophagous insects eventually overcome these defenses and adapt to the plant clade, they sometimes undergo adaptive radiation, resulting in a number of related insect species utilizing related host plant species (Futuyma & Agrawal 2009).

The association between host plant preference and offspring performance is important for understanding how the relationships between plants and phytophagous insects have evolved (Thompson 1988, Nylin *et al*. 1996). In many phytophagous insects, host plants are chosen by the adult females during oviposition (Mayhew 1997, Jones 2022). Female host plant preference can be defined as the hierarchal ordering of different host plant species during oviposition and may reflect host plant qualities such as nutritional value and the occurrence of natural enemies (Thompson 1988, West & Cunningham 2002). Such preference is important for offspring performance, influencing their survival, growth and reproduction. Consequently, selection of suitable sites for oviposition is crucial for offspring fitness and is the only form of parental care shown by many insects (Renwick 1989, Nylin *et al*. 1996, Gamberale-Stille *et al*. 2014, Jones *et al*. 2019). According to the *preference-performance* hypothesis, also known as the *mother knows best* hypothesis (Gripenberg *et al*. 2010), females should therefore evolve to oviposit on host plants on which offspring performance is the highest (Wiklund 1975, Jaenike 1978).

While two meta-analyses have demonstrated that female host plant preference generally is correlated to offspring performance, they also highlight that a many studies have found poor preference-performance correlations (Mayhew 1997, Gripenberg *et al*. 2010). A meta-analysis that separated native and exotic host plant species indicated that the preference-performance hypothesis was applicable to native host plants, but not necessarily to exotic ones (Jones *et al*. 2019). Host plants that are less suitable as food may still be beneficial to utilize if they reduce exposure to natural enemies (Björkman *et al*. 1997) or constitute a source of mutualists (Atsatt 1981), providing alternative explanations for poor preference-performance correlations. In a study on the oviposition of the butterfly *Ogyris amaryllis*, for example, host plants with ant mutualists were preferred over those without ants regardless of plant quality and abundance (Atsatt 1981). Utilizing a more drought resistant host plant species, even if offspring performance is reduced on this species, can also be beneficial (Näsvall *et al*. 2021). Moreover, females do not only choose host plants based on the optimization of their offsprings’ fitness, but also on the optimization of their own fitness (Nylin *et al*. 1996, Scheirs *et al*. 2000). Abundant host plant species associated with an increased oviposition rate may therefore be preferred by females at the cost of individual offspring performance (Nylin *et al*. 1996, Mayhew 1997). This can be seen as examples of a parent-offspring conflict (Gamberale-Stille *et al*. 2014).

Many phytophagous insects utilize different host plants species in different parts of their distribution area, but it is unknown if geographic differences in host plant specialization mainly depend on differences in the availability of host plant species or differences in host plant preference (Fox & Morrow 1981, Thompson 1993). The northern brown argus butterfly (*Aricia artaxerxes*) occurs throughout almost all of Sweden (Figure 1), where it utilizes at least two host plant species – wood cranesbill (*Geranium sylvaticum*) and bloody cranesbill (*G. sanguineum*) (Eliasson *et al*. 2005). Common rockrose (*Helianthemum nummularium*), which is also used by the closely related *A. agestis*, and hoary rockrose (*H. oelandicum*) have also been mentioned as host plants of *A. artaxerxes* in Sweden (Eliasson *et al*. 2005). However, rockroses may be utilized exclusively by the southern lineage *horkei*, which was previously described as a subspecies of *A. artaxerxes* on the Baltic Sea island Öland, but now known to be a hybrid lineage between *A. artaxerxes* and *A. agestis* (Høegh-Guldberg 1974, Eliasson *et al*. 2005, Boman *et al*. 2025). Due to the geographic distribution of its host plant species, it is believed that *A. artaxerxes* utilizes different *Geranium* host plant species in different parts of Sweden, although no studies have yet been made to confirm this (Eliasson *et al*. 2005). In the province of Uppland in central Sweden, where this study was conducted, both *G. sylvaticum* and *G. sanguineum* occur, with the former being more frequent than the latter (Eliasson *et al*. 2005, Jonsell 2010) (Figure 1). The relative usage of the two *Geranium* species by *A. artaxerxes* in Uppland has so far not been investigated in-depth.

**Figure 1.**
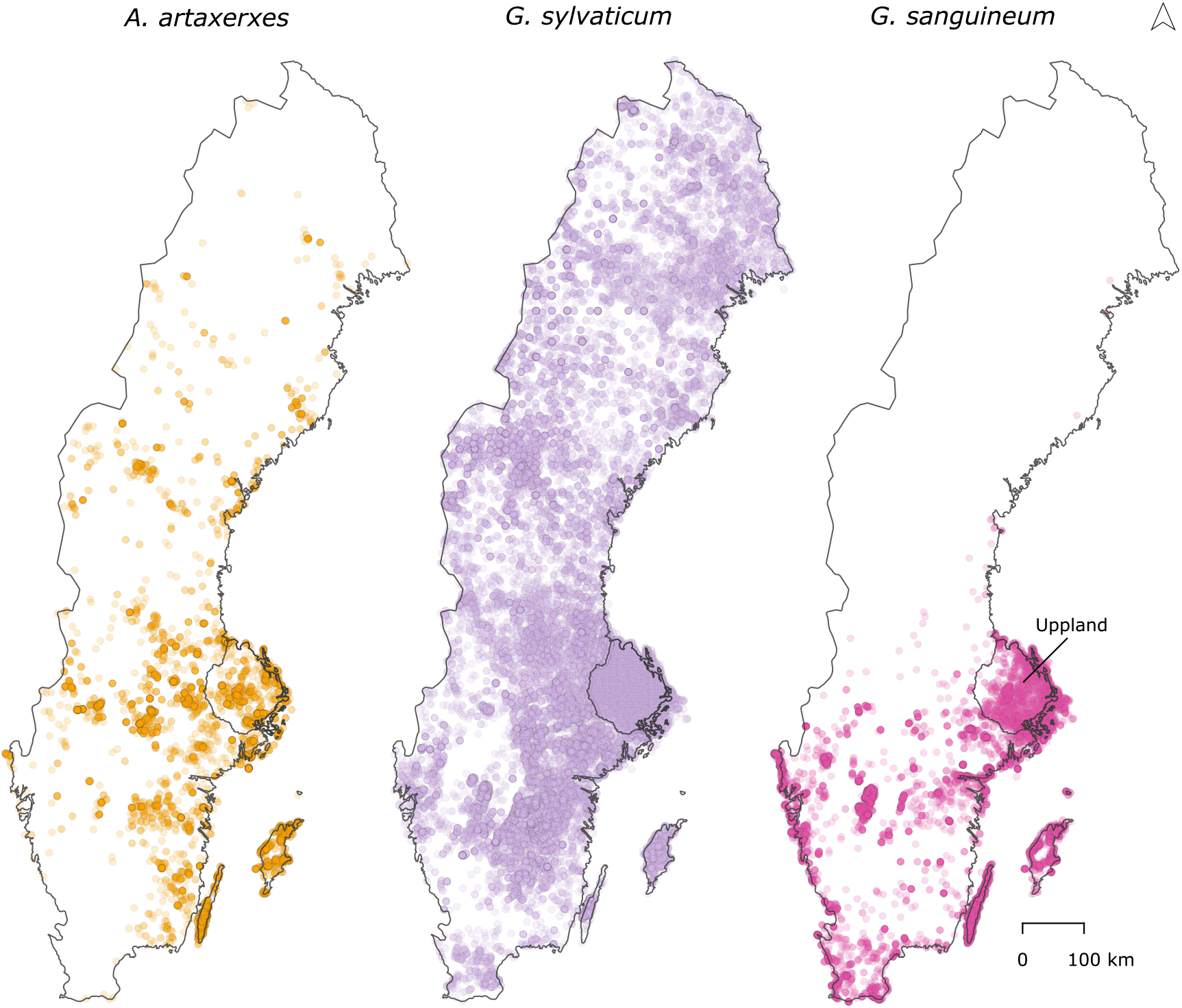
Reported findings of *A. artaxerxes* and the host plants *G. sylvaticum* and *G. sanguineum* in Sweden between year 2000-2025, downloaded from the citizen science database Species *Observation System (Artportalen*) (SLU Swedish Species Information Centre n.d.).

Here, we use *A. artaxerxes* in the province of Uppland, Sweden to investigate 1) if there is a female host plant preference between the most commonly occurring host plant (*G. sylvaticum*) and a rarer host plant (*G. sanguineum*). We also test 2) if there, in according to the preference-performance hypothesis, is a positive connection between female host plant preference and offspring performance. Lastly, we investigate 3) if host plant preference in *A. artaxerxes* is related to the geographic distribution of *A. artaxerxes* and its host plants *G. sylvaticum* and *G. sanguineum* in Uppland.

## Materials and methods

### Host plant preference

For the host plant preference test, *A. artaxerxes* females were collected in the Uppsala area in the province of Uppland, Sweden, between July 2-13, 2024 (localities: 59.8359 N, 17.5880 E; 59.8595 N, 17.5783 E & 59.8394 N, 17.5566 E). The two host plant species (*G. sylvaticum* and *G. sanguineum*) were also collected in the province of Uppland (localities: 59.8359 N, 17.5880 E; 59.8595 N, 17.5783 E; 59.8394 N, 17.5566 E & 60.5093 N, 17.5973 E). During the field work, we observed that both *G. sylvaticum* and *G. sanguineum* were present in the localities where the butterfly females were collected. We placed the collected *A. artaxerxes* females individually in 40 x 40 x 40 cm mesh cages with plastic roofs. We cut the host plants to similar sizes, approximately 30 cm in height, and placed them individually in plastic containers with water. In each cage, we placed four containers with plants, one in each corner with the same host plant species in opposite corners. We switched the placement of the host plant species between the cages. In the center of each cage, we placed a feeding station consisting of *Cirsium* flowers and sugar water. The host plant preference test ran for 72 hours with an 18-hour light + 6-hour darkness regime (light between 3 a.m. and 9 p.m. and dark between 9 p.m. and 3 a.m., which approximately corresponds to conditions in Uppland at the time of collection). After the 72 hours, we counted the number of eggs laid by each female on each host plant species. Out of 28 collected females, 14 survived the 72 hours and 7 of these oviposited.

### Larval growth

During the host plant preference test, 346 eggs were laid by the 7 *A. artaxerxes* females. The eggs were laid individually and not in clusters. The larvae hatching from the eggs were raised on the same host plant as the egg was laid. Host plants were replaced regularly so that larvae always had access to fresh food. If host plants were about to dry up, we placed new plants next to them in the containers. The larvae were raised under the same light conditions used during the host plant preference test (18 hours light + 6 hours darkness). Starting on day 8 after the start of the host plant preference test, we photographed the larvae using a Lumenera Infinity 2-5C microscopy camera mounted on a Leica M165C stereo microscope using the software Infinity analyze version 6.2 (Lumenera corporation 2013). We photographed each cohort of larvae every 3^rd^ day for 12-15 days. For each day and cohort, we photographed up to 20 randomly chosen larvae (5 larvae per plant). To minimize handling, we photographed the larvae on the host plants. We measured the surface area (mm^2^) of the photographed larvae using the polygon selections tool in the software ImageJ version 1.54 (Schneider *et al*. 2012). In total, we took 370 size measurements.

### Geographic distribution

For the geographic distribution, we downloaded data of reported observations of *A. artaxerxes, G. sylvaticum* and *G. sanguineum* in the province of Uppland, Sweden, from the citizen science database Species Observation System *Artportalen* (SLU Swedish Species Information Centre n.d.) on November 27, 2024. For the distribution of *A. artaxerxes*, we used 1.431 observations reported between years 2000-2024. For the distribution *G. sylvaticum* and *G. sanguineum*, we also used data from an inventory of the plants of Uppland (Jonsell 2010). The plant inventory was conducted between year 1991-2005 in 2.742 squares of 2.5 km^2^ covering the province of Uppland. The downloaded data showed in which of these squares *G. sylvaticum* and *G. sanguineum* were noted during the inventory. We created a map of the distribution of *A. artaxerxes, G. sylvaticum* and *G. sanguineum* in Uppland using the 2.5 km^2^ squares from the *plant* inventory in the GIS software QGIS version 3.40 (QGIS Development Team 2024).

### Statistical analyses

The host plant preference, larval growth and geographic distribution were analyzed in the statistical software R version 4.4.1 (R Core Team 2024). Host plant preference was analyzed using a Poisson generalized linear mixed-effect model (GLM) with the number of eggs laid as the response variable and host plant species (*G. sylvaticum* and *G. sanguineum*) as the explanatory variable. The Poisson GLM was chosen because of the count data property of the response variable (the number of eggs laid). Female was used as a random factor to account for the fact that the same female chose between both host plant species, which meant that the samples were not independent. The model was implemented using the *lme4* package, and significances were assessed using the Anova function of the *car* package. Larval growth was analyzed with linear mixed-effects models (LMM) using the *lme4* package (Bates *et al*. 2015), where size (in mm^2^) was the response variable and host plant species (*G. sylvaticum* and *G. sanguineum*) and number of days since oviposition (Day in models) were the explanatory variables. Day^2^ was added to the model to account for non-linear growth. Female (mother) was again used as a random factor since the samples were not independent. The response variable (size, mm^2^) was log transformed since the variance increased with the mean. Since the offspring were kept in groups on the plants where females had laid their eggs, there is a possibility that the density of offspring could affect their growth by competitive interactions. Therefore, in a separate model, we added Density as another factor. Day and Density were scaled to a mean of 0 and a variance of 1, since they were on different scales. Model simplification was performed where non-significant interactions were removed until Akaike information criterion (AIC) stopped decreasing and the model with lowest AIC was chosen (refitted using maximum likelihood, AIC of all models are presented in supplementary table S2 and S4). Although some interaction terms were not individually significant, models including these terms sometimes had lower AIC and were therefore retained to optimize overall model fit. To investigate any difference between the observed and expected distribution of *A. artaxerxes* based on the distribution of the host plants, the number of squares in which *A. artaxerxes, G. sylvaticum* and *G. sanguineum* occurred was analyzed using Pearson’s Chi-squared test. The squares where neither *G. sylvaticum* nor *G. sanguineum* occurred were excluded from the analysis.

## Results

### Host plant preference

We found that the *A. artaxerxes* females laid significantly more eggs on *G. sylvaticum* compared to *G. sanguineum* (Poisson generalized linear mixed-effect model: χ^2^ = 76.623, df = 1, p < 0.001, Figure 2). Of the seven females that laid eggs, five females oviposited on both *G. sylvaticum* and *G. sanguineum* and two females oviposited exclusively on *G. sylvaticum*.

**Figure 2.**
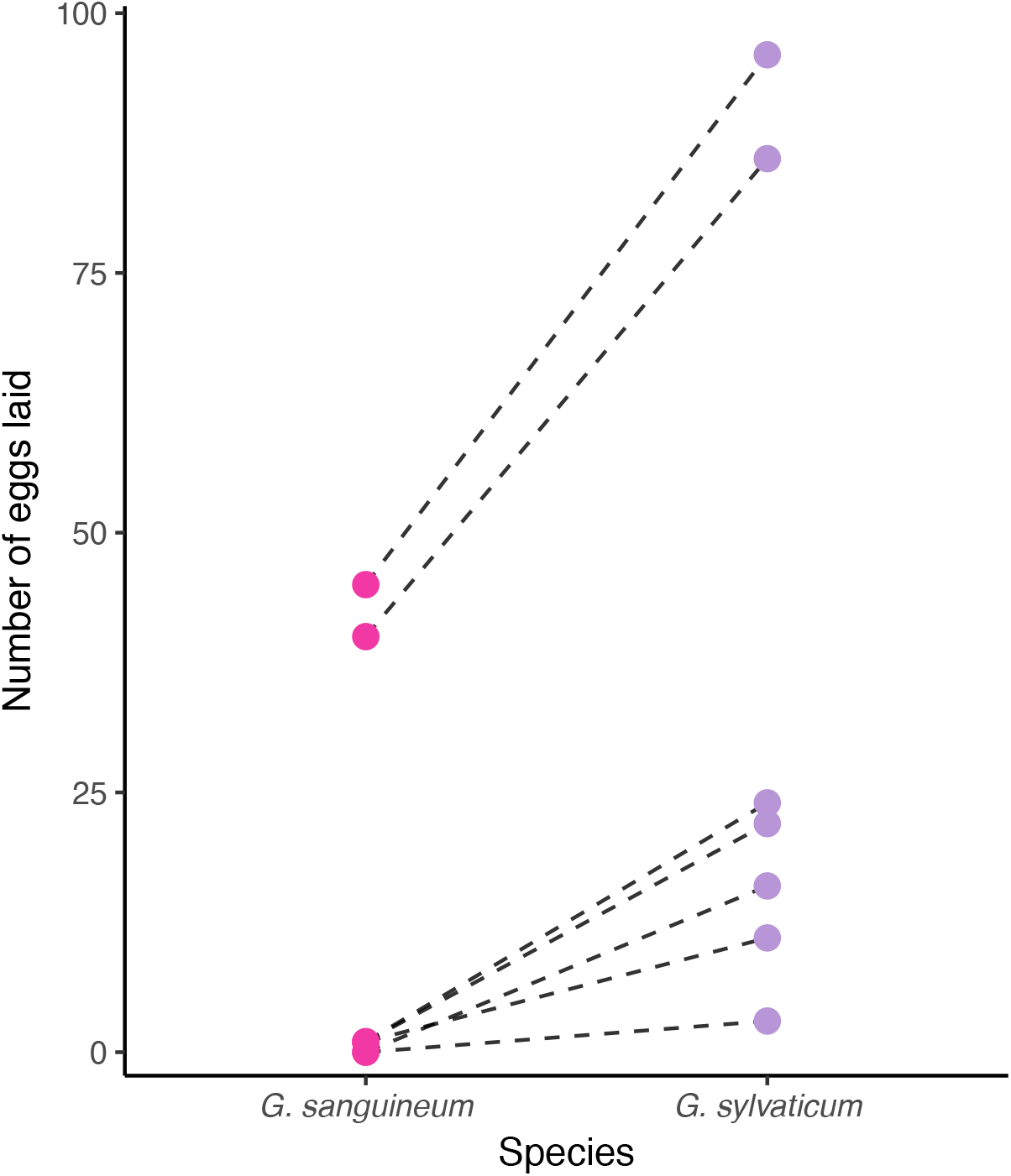
The number of eggs laid by each *A. artaxerxes* female (N = 7) on the host plants *G. sanguineum* and *G. sylvaticum*.

### Larval growth

We found that the growth of the larvae was non-linear (decreasing) as indicated by the significant effect of Day^2^ (Table 1 & Figure 3). Moreover, the significant effect of host plant species demonstrates that the larvae feeding on *G. sylvaticum* were smaller than the larvae feeding on *G. sanguineum* (Table 1, supplementary table 2, Figure 3). There was, however, no interaction between the incline or curvature of the growth between the larvae feeding on *G. sylvaticum* and the larvae feeding on *G. sanguineum* (Supplementary table 1).

**Table 1.** χ^2^, df and p-value of the larval growth linear mixed-effects model with the lowest AIC.

| | $\chi^2$ | df | p |
| --- | --- | --- | --- |
| *Intercept* | 35.87 | 1 | <0.001 |
| *Day* | 858.44 | 1 | <0.001 |
| *Species* | 6.42 | 1 | 0.011 |
| *Day*<sup>2</sup> | 53.04 | 1 | <0.001 |

**Figure 3.**
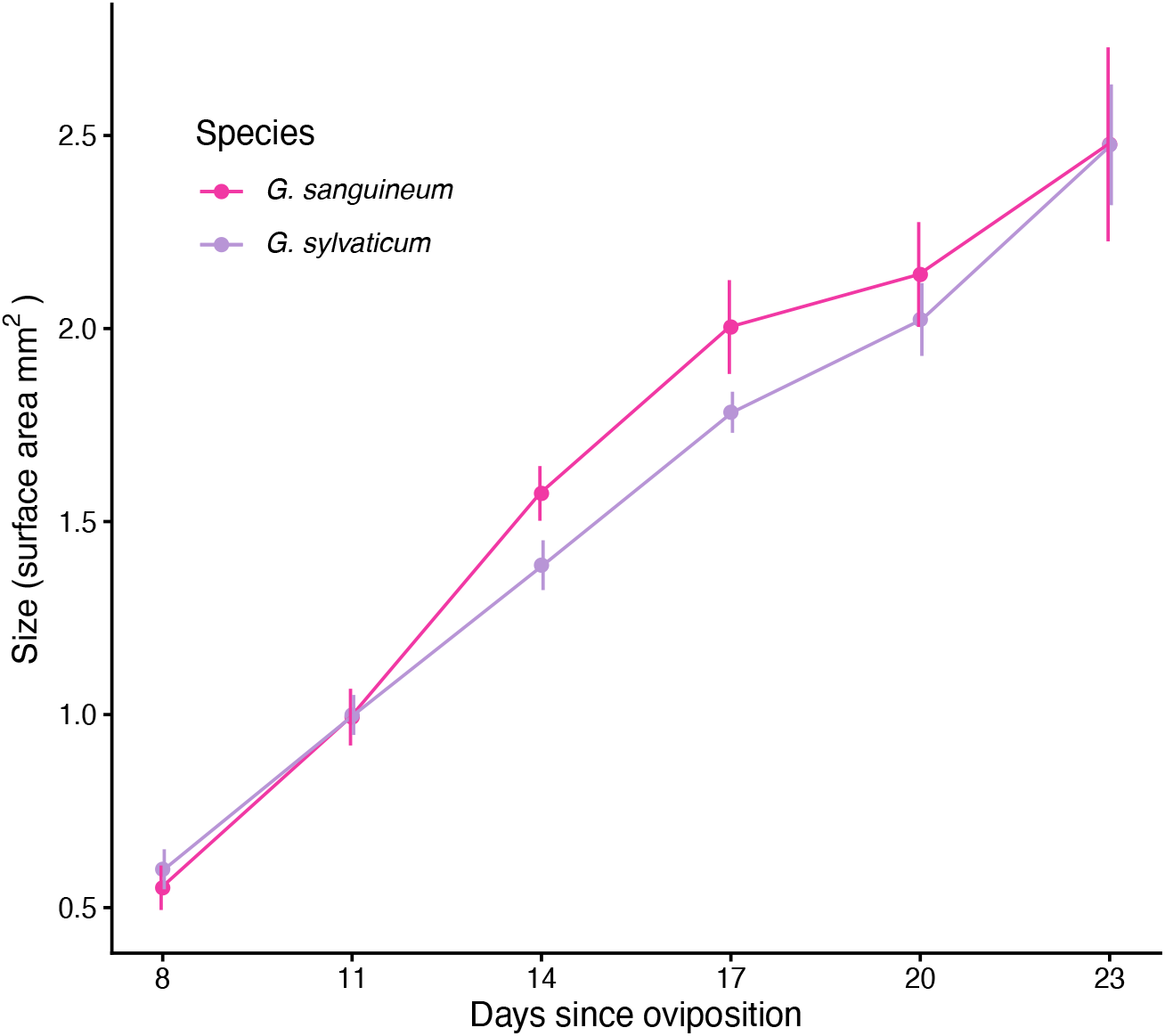
Growth curves of the *A. artaxerxes* larvae feeding on either *G. sylvaticum* or *G. sanguineum* on days 8-23 since oviposition. Dots show mean values and bars represent standard errors.

To investigate the possibility that any growth differences were driven by competition, we fitted a separate set of models that also included the density of larvae. After model simplification (supplementary table 3), we found qualitatively similar results, where larval size was smaller for offsprings feeding on *G. sylvaticum*. Larval density did not have any direct effect on size, nor was it included in any significant interactions, although model selection indicated that it’s inclusion in the final model improved model fit (Table 2, supplementary table 4).

**Table 2.** χ^2^, df and p-value of the larval growth linear mixed-effects model with the lowest AIC, including the effect of larval density.

| | $\chi^2$ | df | p |
| --- | --- | --- | --- |
| *Intercept* | 29.05 | 1 | <0.001 |
| *Day* | 266.72 | 1 | <0.001 |
| *Species* | 4.03 | 1 | 0.044 |
| *Density* | 2.65 | 1 | 0.103 |
| *Day*<sup>2</sup> | 29.61 | 1 | <0.001 |
| *Day* × *Species* | 0.60 | 1 | 0.441 |
| *Day* × *Density* | 0.01 | 1 | 0.929 |
| *Species* × *Density* | 2.25 | 1 | 0.134 |
| *Species* × *Day*<sup>2</sup> | 2.41 | 1 | 0.121 |
| *Density* × *Day*<sup>2</sup> | 1.99 | 1 | 0.159 |
| *Day* × *Species* × *Density* | 2.87 | 1 | 0.090 |

### Geographic distribution

*G. sylvaticum* and *G. sanguineum* occurred in 96% and 47% of the 2387 2.5 km^2^ squares respectively (Figure 4). *A. artaxerxes* commonly occurred where only *G. sylvaticum* was present, (in 40% of the squares) but rarely occurred where only *G. sanguineum* was present (in 1% of the squares). However, we found that there was a significant difference between the observed and expected distribution of *A. artaxerxes* based on the distribution of *G. sylvaticum* and *G. sanguineum* (Pearson’s Chi-squared test: χ^2^ = 25.973, df = 2, p < 0.001, Figure 5). *A. artaxerxes* mainly (in 55% of the squares where it was present) occurred in squares where both *G. sylvaticum* and *G. sanguineum* were present, and more frequently so than expected (Figure 4 & Figure 5). However, *A. artaxerxes* occurred less frequently than expected where only *G. sylvaticum* or only *G. sanguineum* was present (Figure 5).

**Figure 4.**
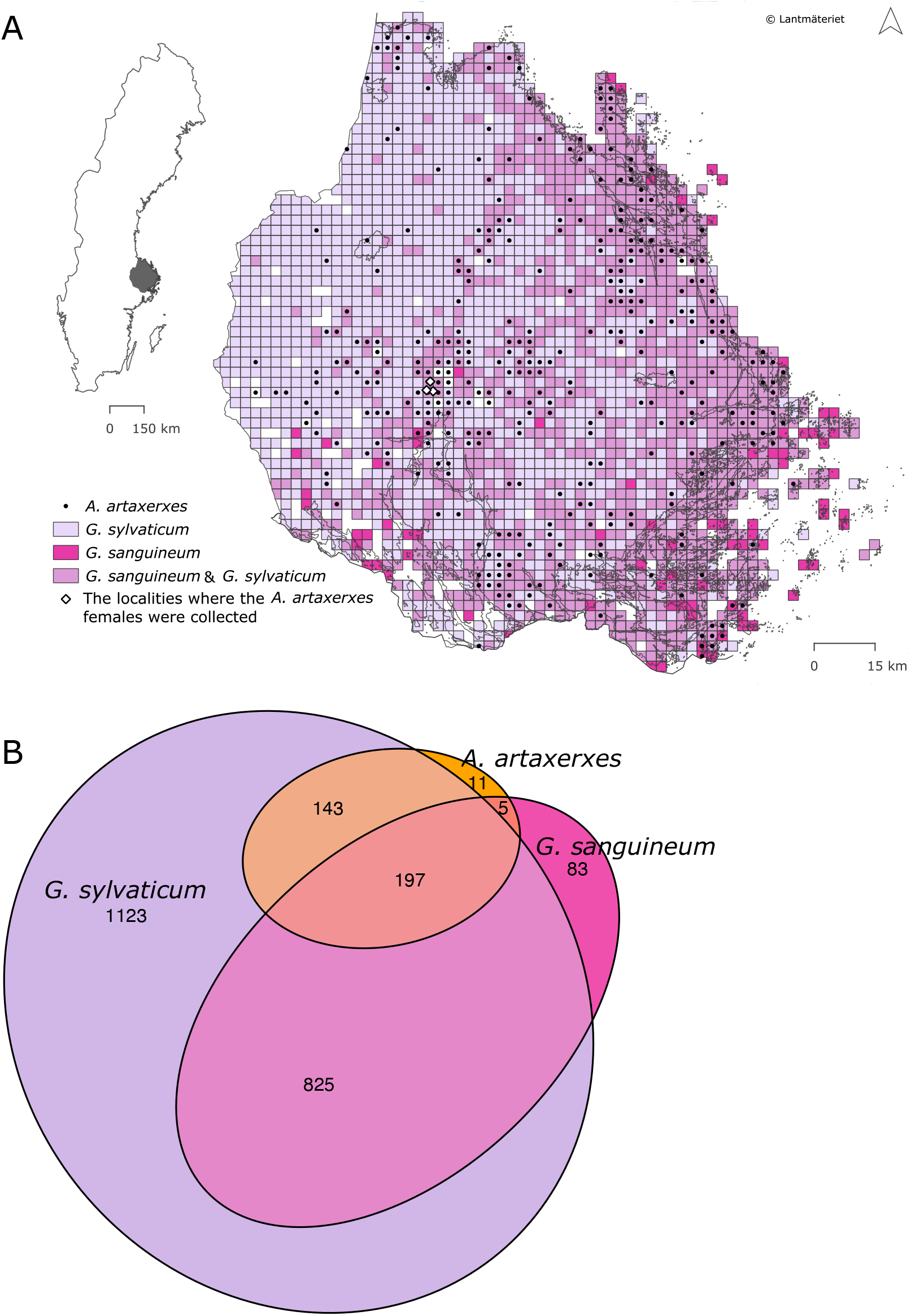
A) The distribution of *A. artaxerxes, G. sylvaticum* and *G. sanguineum* in the province of Uppland, Sweden, in 2.5 km2 squares as well as the localities where the *A. artaxerxes* females were collected for the host plant preference test. The map was generated using a basemap from Lantmäteriet. B) Venn diagram of the number of 2.5 km2 squares in Uppland in which *A. artaxerxes, G. sylvaticum* and *G. sanguineum* occurred and their overlap.

**Figure 5.**
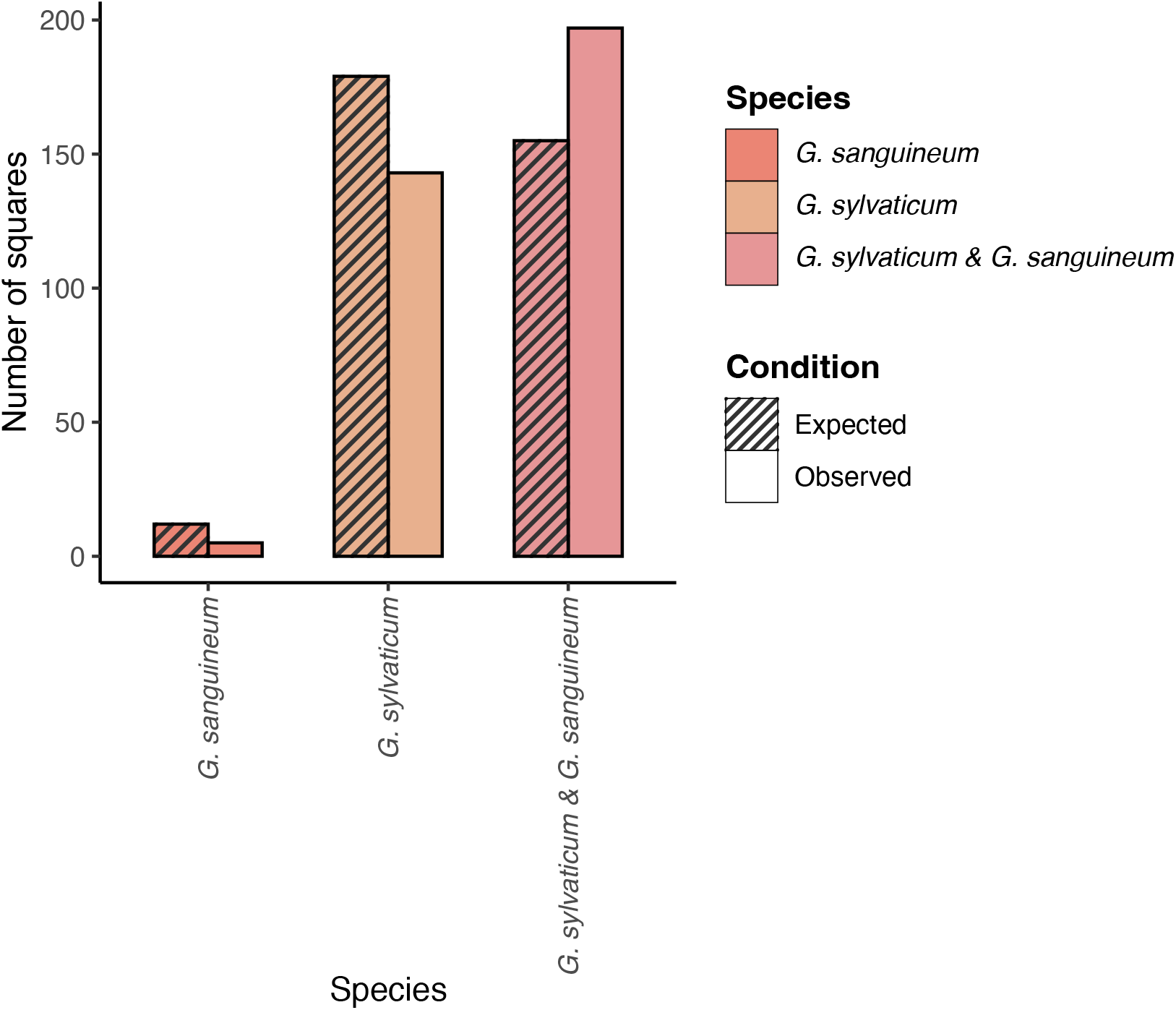
Expected and observed number of 2.5 km^2^ squares in the province of Uppland, Sweden, where *A. artaxerxes* occurred and where either only *G. sanguineum*, only *G. sylvaticum* or both *G. sylvaticum* and *G. sanguineum* were present.

## Discussion

### Preference and performance

Contrary to the preference-performance hypothesis (Wiklund 1975, Jaenike 1978), we found that the *A. artaxerxes* females from the Uppsala area in the province of Uppland, Sweden, strongly prefer ovipositing on *G. sylvaticum* rather than on *G. sanguineum*, even though the larvae feeding on *G. sylvaticum* grew slightly smaller than those feeding on *G. sanguineum*. This result held true even when accounting for the higher larval density and possible stronger competitive interactions on *G. sylvaticum*. Our results indicate that there are other factors than offspring performance influencing the *A. artaxerxes* females’ host plant preference. Even though female host plant preference generally is connected to offspring performance (Mayhew 1997, Gripenberg *et al*. 2010, Jones *et al*. 2019, Zanco *et al*. 2025), many exceptions to the preference-performance hypothesis have been documented (Wiklund 1975, Chew 1977, Courtney 1981, Valladares & Lawton 1991, Ohsaki & Sato 1994, Underwood 1994, Björkman *et al*. 1997, Berdegué *et al*. 1998, Clark *et al*. 2011, Davis & Cipollini 2014, Näsvall *et al*. 2021), including this study. However, it may still be adaptive for females not to choose the host plant with the highest quality as food for the offspring (Gripenberg *et al*. 2010). In the green-veined white (*Pieris napi*) (Ohsaki & Sato 1994) and the redheaded pine sawfly (*Neodiprion sertifer*) (Björkman *et al*. 1997), oviposition on less nutritious host plants may reduce exposure to parasites. In the amaryllis azure (*Ogyris amaryllis*), oviposition on a less nutritious host plant species can provide a source of ant mutualists (Atsatt 1981). In the common wood white (*Leptidea sinapis*), females may prefer ovipositing on a less nutritious host plant species due to it being more drought resistant (Näsvall *et al*. 2021). Sometimes, however, female host plant preference does not seem to be adaptive, for example when females oviposit on exotic host plant species with low offspring survival, such as for example documented in the old world swallowtail (*Papilio machaon*) (Wiklund 1975), the green-veined white (*Pieris napi*) (Chew 1977) and the West Virginia white (*Pieris virginiensis*) (Davis & Cipollini 2014). In this way, exotic plants can negatively impact native butterfly species (Graves & Shapiro 2003), and an explanation for this behavior is that adaptations have not yet evolved for females to avoid ovipositing on exotic species (Wiklund 1975, Chew 1977).

Another factor influencing female host plant preference is differences in abundance between host plant species, as females are more likely to oviposit on lower quality host plant species if they are more abundant than higher quality host plant species (Rausher 1980, Singer 1983, Nylin *et al*. 1996, Mayhew 1997, West & Cunningham 2002). Females that search preferentially for a more common host plant species may discover more plants and lay more eggs during their lifetimes compared to females that search preferentially for a rarer host plant species, thereby increasing their fitness (Rausher 1980, Nylin *et al*. 1996). Abundant host plant species may therefore, even at the cost of individual offspring performance, be preferred by females and trade-offs may be made between quality and quantity of offspring (Nylin *et al*. 1996, Mayhew 1997). This may be an important factor shaping the *A. artaxerxes* females’ preference for *G. sylvaticum* since *G. sylvaticum* is more commonly occurring than *G. sanguineum* in Uppland, both overall and in the areas where *A. artaxerxes* occurs. Further, larvae feeding on *G. sylvaticum* are smaller than larvae feeding on *G. sanguineum*, but only slightly. The potential difference in female fitness based on host plant abundance (Rausher 1980, Nylin *et al*. 1996) and relatively low difference in offspring performance may explain why the *A. artaxerxes* females, contrary to the preference-performance hypothesis (Wiklund 1975, Jaenike 1978), do not prefer to oviposit on the host plant species on which offspring performance is the highest. If the locally most common host plant also offers best offspring performance, this plant should be preferred. This has recently been demonstrated in the closely related Brown Argus (*A. agestis*) in the UK, where recently established local populations of this expanding species have switched their oviposition preference from the ancestral perennial Rockrose (*Helianthemum nummularium*) to the more widespread *Geranium molle*, where offspring performance is also higher (Zanco *et al*. 225). Negative preference-performance correlations, such as the one found in *A. artaxerxes* in our study, may indicate that there is a conflict between female fitness and direct offspring fitness, i.e. a parent-offspring conflict, where females maximizing their fitness by ovipositing on suboptimal host plants at the expense of offspring performance (Gamberale-Stille *et al*. 2014).

### The role of host plant distribution

When investigating the geographical distribution, we found that *A. artaxerxes* mainly occurs in areas where both *G. sylvaticum* and *G. sanguineum* occur, and significantly more frequently so than expected based on the distribution of the two host plants. *A. artaxerxes* also commonly occurs in areas where only *G. sylvaticum* occurs and, although very rarely, where only *G. sanguineum* occurs. It should however be noted that the availability of host plants is likely not the only factor that determines the realized niche of *A. artaxerxes*, and its common occurrence in areas where both host plant species occur may reflect suitable microclimate and abiotic conditions. Although these results are based on a relatively high number of citizen-reported *A. artaxerxes* observations (1431 observations) (SLU Swedish Species Information Centre n.d.), the observations are patchily distributed, and the actual distribution of *A. artaxerxes* probably exceeds the distribution as reported in the database, which may affect the results. In contrast, the distributions of host plants is based on atlas inventories (Jonsell 2010) and should be considered robust. Still, there are areas where only one host plant species occurs, indicating that there are areas where either *G. sanguineum* or *G. sylvaticum* cannot be used. Regarding areas where both host plant species occur, the host plant preference test indicates that *A. artaxerxes* mainly utilizes *G. sylvaticum* in these areas, but that *G. sanguineum* probably is utilized to a smaller degree as well. In partial agreement with the previous estimation (Eliasson *et al*. 2005), these results indicate that *A. artaxerxes* probably mainly, but not exclusively, utilizes *G. sylvaticum* in Uppland. The results also indicate that the *A. artaxerxes* females prefer the host plant species that is most commonly occurring in Uppland. Studies of phytophagous insects using different host plant species in different parts of their distribution area show that some populations have diverged in host plant preference (Gotthard *et al*. 2004, Näsvall *et al*. 2021) while others have not (Jaenike 1989, Thompson 1993). The *A. artaxerxes* females’ preference for *G. sylvaticum* may therefore be a local adaptation in host plant preference, although studying *A. artaxerxes’* host plant preference in other geographical areas where other host plant species are utilized would be necessary to determine this.

### Conclusions

Contrary to the preference-performance hypothesis (Wiklund 1975, Jaenike 1978), we found that the *A. artaxerxes* females from the province of Uppland, Sweden, prefer ovipositing on *G. sylvaticum* compared to *G. sanguineum*, even though larvae feeding on *G. sylvaticum* are slightly smaller than those feeding on *G. sanguineum*. Since *G. sylvaticum* is more commonly occurring than *G. sanguineum* in Uppland, potential differences in female fitness based on host plant abundance (Rausher 1980, Nylin *et al*. 1996) and low difference in offspring performance may explain why the *A. artaxerxes* females do not prefer to oviposit on the host plant species on which offspring performance is the highest. *G. sylvaticum* being the most commonly occurring and probably most utilized host plant in Uppland indicates that the *A. artaxerxes* females’ preference for *G. sylvaticum* could be a local adaptation in host plant preference, although further studies are needed to determine this. Overall, the results show that factors other than offspring performance, such as geographic distribution and commonness of the host plants, may influence female host plant preference in *A. artaxerxes*.

## Author contributions

Vanda Larsson Åberg (Conceptualization [supporting], Data curation [lead], Investigation [lead], Methodology [supporting], Formal analysis [supporting], Visualization [equal], Writing – original draft [lead], Writing – review & editing [equal]), Jesper Boman (Visualization [supporting], Writing – review & editing [equal]), Niclas Backström (Conceptualization [equal], Funding acquisition [equal], Methodology [lead], Resources [equal], Supervision [supporting], Writing – review & editing [equal]), Martin I. Lind (Conceptualization [equal], Formal analysis [lead], Funding acquisition [equal], Resources [equal], Supervision [lead], Visualization [equal], Writing – original draft [supporting], Writing – review & editing [equal]).

## References

Abrams PA. 2000. The evolution of predator-prey interactions: Theory and evidence. Annual Review of Ecology and Systematics 31: 79–105.

Anderson RM, May RM. 1982. Coevolution of hosts and parasites. Parasitology 85: 411–426.

Atsatt PR. 1981. Ant-dependent food plant selection by the mistletoe butterfly Ogyris amaryllis (Lycaenidae). Oecologia 48: 60–63.

Bates D, Mächler M, Bolker B, Walker S. 2015. Fitting linear mixed-effects models using lme4. Journal of Statistical Software 67: 1–48.

Bennett RN, Wallsgrove RM. 1994. Secondary metabolites in plant defence mechanisms. New Phytologist 127: 617–633.

Berdegué M, Reitz SR, Trumble JT. 1998. Host plant selection and development in Spodoptera exigua: do mother and offspring know best? Entomologia Experimentalis et Applicata 89: 57–64.

Bernays E, Graham M. 1988. On the evolution of host specificity in phytophagous arthropods. Ecology 69: 886–892.

Björkman C, Larsson S, Bommarco R. 1997. Oviposition preferences in pine sawflies: A trade-off between larval growth and defence against natural enemies. Oikos 79: 45–52.

Boman J, Nolen ZJ, Backström N. 2025. On the origin of an insular hybrid butterfly lineage. Evolution 79: 510–524.

Bronstein JL, Alarcón R, Geber M. 2006. The evolution of plant–insect mutualisms. New Phytologist 172: 412–428.

Chew FS. 1977. Coevolution of pierid butterflies and their cruciferous foodplants. II. The distribution of eggs on potential foodplants. Evolution 31: 568–579.

Clark KE, Hartley SE, Johnson SN. 2011. Does mother know best? The preference– performance hypothesis and parent–offspring conflict in aboveground– belowground herbivore life cycles. Ecological Entomology 36: 117–124.

Connell JH. 1980. Diversity and the coevolution of competitors, or the ghost of competition past. Oikos 35: 131–138.

Courtney SP. 1981. Coevolution of pierid butterflies and their cruciferous foodplants. Oecologia 51: 91–96.

Davis SL, Cipollini D. 2014. Do mothers always know best? Oviposition mistakes and resulting larval failure of Pieris virginiensis on Alliaria petiolata, a novel, toxic host. Biological Invasions 16: 1941–1950.

Ehrlich PR, Raven PH. 1964. Butterflies and plants: A study in coevolution. Evolution 18: 586–608.

Eliasson CU, Ryrholm N, Gärdenfors U. 2005. Nationalnyckeln till Sveriges flora och fauna: Fjärilar: Dagfjärilar. Hesperiidae – Nymphalidae. Artdatabanken, Sveriges lantbruksuniversitet, Uppsala.

Futuyma DJ, Agrawal AA. 2009. Macroevolution and the biological diversity of plants and herbivores. Proceedings of the National Academy of Sciences 106: 18054–18061.

Gamberale-Stille G, Söderlind L, Janz N, Nylin S. 2014. Host plant choice in the comma butterfly–larval choosiness may ameliorate effects of indiscriminate oviposition. Insect Science 21: 499–506.

Gotthard K, Margraf N, Rahier M. 2004. Geographic variation in oviposition choice of a leaf beetle: the relationship between host plant ranking, specificity, and motivation. Entomologia Experimentalis et Applicata 110: 217–224.

Graves SD, Shapiro AM. 2003. Exotics as host plants of the California butterfly fauna. Biological Conservation 110: 413–433.

Gripenberg S, Mayhew PJ, Parnell M, Roslin T. 2010. A meta-analysis of preference– performance relationships in phytophagous insects. Ecology Letters 13: 383–393.

Høegh-Guldberg O. 1974. Aricia artaxerxes F. ssp. horkei H.-Guld.: (Lep.Rhopalocera). Descriptionof the preliminary stages and a crossing with A.a.ssp.rambringi H.-Guld. Aricia Studies No. 13. Insect Systematics & Evolution 4: 225–232.

IUCN. 2025. The IUCN Red List of Threatened Species. WWW-dokument 2025-: https://www.iucnredlist.org/en. Hämtad 2025-05-19.

Jaenike J. 1978. On optimal oviposition behavior in phytophagous insects. Theoretical population biology 14: 350–356.

Jaenike J. 1989. Genetic population structure of Drosophila tripunctata: Patterns of variation and covariation of traits affecting resource use. Evolution 43: 1467–1482.

Jones LC. 2022. Insects allocate eggs adaptively according to plant age, stress, disease or damage. Proceedings of the Royal Society B: Biological Sciences 289: 20220831.

Jones LC, Rafter MA, Walter GH. 2019. Insects allocate eggs adaptively across their native host plants. Arthropod-Plant Interactions 13: 181–191.

Jonsell L. 2010. Upplands flora. SBF-förlaget, Uppsala.

Lumenera corporation. 2013. Infinity analyze.

Mayhew PJ. 1997. Adaptive patterns of host-plant selection by phytophagous insects. Oikos 79: 417–428.

Nylin S, Janz N, Wedell N. 1996. Oviposition plant preference and offspring performance in the comma butterfly: Correlations and conflicts. Entomologia Experimentalis et Applicata 80: 141–144.

Näsvall K, Wiklund C, Mrazek V, Künstner A, Talla V, Busch H, Vila R, Backström N. 2021. Host plant diet affects growth and induces altered gene expression and microbiome composition in the wood white (Leptidea sinapis) butterfly. Molecular Ecology 30: 499–516.

Ohsaki N, Sato Y. 1994. Food plant choice of pieris butterflies as a trade-off between parasitoid avoidance and quality of plants. Ecology 75: 59–68.

QGIS Development Team. 2024. QGIS Geographic Information System.

R Core Team. 2024. R: A language and environment for statistical computing.

Rausher MD. 1980. Host abundance, juvenile survival, and oviposition preference in Battus philenor. Evolution 34: 342–355.

Renwick JAA. 1989. Chemical ecology of oviposition in phytophagous insects. Experientia 45: 223–228.

Scheirs J, De Bruyn L, Verhagen R. 2000. Optimization of adult performance determines host choice in a grass miner. Proceedings: Biological Sciences 267: 2065–2069.

Schneider CA, Rasband WS, Eliceiri KW. 2012. NIH Image to ImageJ: 25 years of image analysis. Nature Methods 9: 671–675.

Singer MC. 1983. Determinants of multiple host use by a phytophagous insect population. Evolution 389–403.

SLU Swedish Species Information Centre. n.d. Artportalen. https://artfakta.se/. Accessed May 4, 2025.

Thompson JN. 1989. Concepts of coevolution. Trends in Ecology & Evolution 4: 179–183.

Thompson JN. 1993. Preference hierarchies and the origin of geographic specialization in host use in swallowtail butterflies. Evolution 47: 1585–1594.

Thompson JN. 1988. Evolutionary ecology of the relationship between oviposition preference and performance of offspring in phytophagous insects. Entomologia Experimentalis et Applicata 47: 3–14.

Underwood DLA. 1994. Intraspecific variability in host plant quality and ovipositionaI preferences in Eucheira socialis (Lepidoptera: Pieridae). Ecological Entomology 19: 245–256.

Valladares G, Lawton JH. 1991. Host-plant selection in the holly leaf-miner: Does mother know best? Journal of Animal Ecology 60: 227–240.

West SA, Cunningham PJ. 2002. A general model for host plant selection in phytophagous insects. Journal of Theoretical Biology 214: 499–513.

Wiklund C. 1975. The evolutionary relationship between adult oviposition preferences and larval host plant range in Papilio machaon L. Oecologia 18: 185–197.

Zanco B, Jong M de, Widman E, Camus FM, Bridle J. 2025. Does mother know best? Range-wide narrowing of host preference in Aricia agestis confers fitness benefits but may incur long-term costs. 2025.09.04.674201.

