## Supplementary materials for "No preference for performance: Host plant preference, offspring performance and host plant distribution in the butterfly *Aricia artaxerxes*"

**Supplementary table 1.** Model structure and Akaike information criterion (AIC) of the larval growth linear mixed-effects models.

| *Model structure* | *AIC* |
|---|---|
| Day + $Day^2$ + Species + Species:Day + Species:$Day^2$ | 269.44 |
| Day + $Day^2$ + Species + Species:Day | 269.24 |
| Day + $Day^2$ + Species + Species:$Day^2$ | 267.81 |
| Day + $Day^2$ + Species | 267.45 |

**Supplementary table 2.** Parameter estimates (mean SE) of the fixed effects in the model with lowest AIC (Day + $Day^2$ + Species).

| **Parameter** | **Mean** | **SE** |
|---|---|---|
| Intercept | 0.550 | 0.092 |
| Day | 0.521 | 0.018 |
| Species | -0.108 | 0.043 |
| $Day^2$ | -0.138 | 0.019 |

**Supplementary table 3.** Model structure and Akaike information criterion (AIC) of the larval growth linear mixed-effects models including larval density.

| *Model structure* | *AIC* |
|---|---|
| Day + Species + Density + $Day^2$ + Day:Species + Day:Density + Species:Density + Species:$Day^2$ + Density:$Day^2$ + Day:Species:Density + $Day^2$:Species:Density | 257.13 |
| Day + Species + Density + $Day^2$ + Day:Species + Day:Density + Species:Density + Species:$Day^2$ + Density:$Day^2$ + $Day^2$:Species:Density | 258.06 |
| Day + Species + Density + $Day^2$ + Day:Species + Day:Density + Species:Density + Species:$Day^2$ + Density:$Day^2$ + Day:Species:Density | 255.15 |
| Day + Species + Density + $Day^2$ + Day:Species + Day:Density + Species:Density + Species:$Day^2$ + Density:$Day^2$ | 256.09 |
| Day + Species + Density + $Day^2$ + Day:Species + Species:Density + Species:$Day^2$ + Density:$Day^2$ | 262.92 |
| Day + Species + Density + $Day^2$ + Day:Species + Day:Density + Species:Density + Species:$Day^2$ | 256.15 |
| Day + Species + Density + $Day^2$ + Day:Density + Species:Density + Species:$Day^2$ + Density:$Day^2$ | 255.44 |
| Day + Species + Density + $Day^2$ + Day:Species + Day:Density + Species:Density + Density:$Day^2$ | 256.42 |
| Day + Species + Density + $Day^2$ + Day:Species + Day:Density + Species:$Day^2$ + Density:$Day^2$ | 256.66 |

**Supplementary table 4.** Parameter estimates (mean SE) of the fixed effects in the model with lowest AIC that includes Density.

| Parameter | Mean | SE |
|---|---|---|
| Intercept | 0.556 | 0.103 |
| Day | 0.537 | 0.033 |
| Species | -0.152 | 0.076 |
| Density | 0.070 | 0.043 |
| $Day^2$ | -0.185 | 0.034 |
| Day : Species | -0.030 | 0.039 |
| Day : Density | -0.003 | 0.038 |
| Species : Density | -0.078 | 0.052 |
| Species : $Day^2$ | 0.063 | 0.041 |
| Density : $Day^2$ | -0.028 | 0.020 |
| Day : Species : Density | 0.073 | 0.043 |
